# ATM-deficient lung, prostate and pancreatic cancer cells are acutely sensitive to the combination of olaparib and the ATR inhibitor AZD6738

**DOI:** 10.1101/2020.03.13.991166

**Authors:** Nicholas R. Jette, Suraj Radhamani, Ruiqiong Ye, Yaping Yu, Greydon Arthur, Siddhartha Goutam, Tarek A Bizmar, Mehul Kumar, Pinaki Bose, Steven Yip, Michael Kolinsky, Susan P. Lees-Miller

## Abstract

**Background:** The ataxia telangiectasia mutated (ATM) protein kinase is mutated in several human cancers, presenting potential opportunities for targeted cancer therapy. We previously reported that the poly-ADP ribose polymerase (PARP) inhibitor olaparib induced transient G2 arrest but not cell death in ATM-deficient A549 lung cancer cells, while the combination of olaparib with the ATM-, Rad3-related (ATR) inhibitor VE-821 induced cell death. Here, we show that the clinically relevant ATR inhibitor, AZD6738, sensitizes ATM-deficient A549 lung, prostate and pancreatic cancer cells to olaparib.

**Methods:** ATM was depleted from A549 lung cancer cells, PC-3 prostate cancer cells and Panc 10.05 pancreatic cancer cells, and the effects of olaparib alone and in combination with AZD6738 were determined.

**Results:** The combination of olaparib plus AZD6738 induced cell death in ATM-deficient lung, prostate and pancreatic cancer cells with little effect on their ATM-proficient counterparts.

**Conclusions:** Lung, prostate and pancreatic patients whose tumours exhibit loss or inactivation of ATM may benefit from combination of a PARP inhibitor plus an ATR inhibitor.

## Introduction

A goal of precision oncology is to achieve better tumour control by targeting tumours based on their specific genetic characteristics. Perhaps one of the best examples of this approach is the development of poly-ADP ribose polymerase (PARP) inhibitors to target tumours with defects in DNA damage response (DDR) genes ^1^. PARP inhibitors were first shown to target breast cancer cells with depletion of the breast and ovarian cancer susceptibility proteins BRCA1 and BRCA2 ^2,3^ and are now FDA-approved for use in *BRCA*-deficient ovarian cancer ^4^ with clinical trials showing potential in *BRCA*-deficient breast, prostate and pancreatic cancers ^5-12^. BRCA1 and 2 play important roles in the detection and repair of DNA double strand breaks (DSBs) through the Homologous Recombination (HR) repair pathway ^13^, leading to the idea that cells with deficiency in DSB repair pathways may be sensitive to PARP inhibitors. Indeed, siRNA depletion of other proteins involved in the cellular response to DSBs, including Ataxia Telangiectasia Mutated (ATM), also sensitized cancer cells to PARP inhibition ^14,15^.

ATM is an apex signalling kinase that plays a central role in the cellular response to DNA damage ^16^. Loss of both copies of *ATM* leads to ataxia telangiectasia (A-T), a condition characterized by neurological degeneration, cancer predisposition and immune deficiencies ^17^. *ATM* is mutated in a variety of human cancers (reviewed in ^18^), leading us to hypothesize that PARP inhibitors could have therapeutic potential in cancers with loss or mutation of *ATM*. Indeed, up to 40% of mantle cell lymphoma (MCL) have loss of *ATM* ^19^, and we showed that ATM-deficient MCL cells are sensitive to the PARP inhibitor olaparib in both cell line and animal models ^20^. Similarly, ATM-deficient Chronic Lymphocytic Leukemia (CLL) cells are sensitive to PARP inhibitors ^21^. PARP inhibitor sensitivity in ATM-deficient MCL cells was enhanced when p53 was also deleted or inactivated, and p53-deficient MCL cell lines were sensitive to PARP inhibitors in the presence of an ATM kinase inhibitor, KU55933 ^22^. *ATM* has also been reported to be mutated in ∼12.5% of gastric cancers ^23,24^ and p53-deficient gastric cancer cell lines with depletion of ATM, or in which ATM is inhibited using KU55933, are also sensitive to olaparib ^25^. Similarly, ATM is mutated in approximately 16% of colorectal cancers, ATM-deficient colorectal cancer cell lines are sensitive to olaparib and this sensitivity was enhanced when *TP53* was also deleted ^26^.

Recently, we showed that olaparib causes reduced proliferation and reversible G2 arrest in ATM-deficient p53-proficient A549 lung cancer cells, but not cell death ^27^. However, cell death was induced when olaparib-treated, ATM-deficient cells were incubated with the ATR inhibitor VE-821. Thus, our results suggested that in ATM-deficient A549 cells, olaparib alone is cytostatic and the combination of olaparib with an ATR inhibitor is required to induce cell death ^27^. In this study, we examined whether ATM-deficient cells from additional cancer types, specifically prostate and pancreatic cancers, were also sensitive to the combination of olaparib and an ATR inhibitor. Instead of VE-821, a compound widely used in laboratory research ^28^, we used AZD6738, an ATR inhibitor in clinical trials for a number of solid tumours ^29,30^.

Prostate cancer is the most commonly diagnosed cancer in men in Canada ^31^, accounting for almost 10% of all cancer deaths amongst males. Next generation sequencing has indicated that up to 10% of prostate cancers harbour mutations in at least one DDR gene, with *BRCA2* and *ATM* being the most commonly mutated ^32^. Moreover, the results of the recent TOPARP-B clinical trial support a role for olaparib in metastatic castration-resistant prostate cancer with DDR gene aberrations ^8^.

Pancreatic cancer is one of the most lethal forms of cancer with a 10-year survival of less than 10%, and accounts for approximately 6% of all cancer deaths in Canadians ^31^. *ATM* is mutated in some hereditary forms of pancreatic cancer ^33,34^ as well as ∼ 6% of sporadic pancreatic adenocarcinoma, with 4% of overall mutations resulting in deletion of the ATM protein ^35^. In patients with metastatic pancreatic cancer and germline BRCA1/2 mutations, olaparib maintenance treatment following first-line platinum-based chemotherapy resulted in a significant improvement in progression-free survival, in contrast to placebo ^7^.

The encouraging evidence of clinical efficacy of therapy targeting the heterogeneous group of DDR alterations in these tumor groups ^7-12^ prompted us to ask whether prostate and pancreatic cancer cell lines with ATM deficiency would be sensitive to PARP inhibitor either alone, or in combination with AZD6738.

## Materials and methods

### Cell lines

A549 lung adenocarcinoma, Panc 10.05 pancreatic cancer and PC-3 prostate cancer cell lines were obtained from ATCC and checked regularly for mycoplasma. Cells were maintained in a humidified incubator in the presence of 5% CO_2_ at 37°C in either Dulbecco’s Modified Eagle Medium (DMEM) (ThermoFisher Scientific, MA, USA) plus 10% (w/v) Hyclone Fetalclone III Serum (A549 cells), F-12K nutrient mixture (1X) Kaighn’s modification (ThermoFisher Scientific, MA, USA) with 10% (w/v) Fetal Bovine Serum (FBS) (PC-3), or RPMI 1649 (1X) (ThermoFisher Scientific, MA, USA) in the presence of 15% (w/v) Hyclone Fetalclone III Serum (Panc 10.05 cells). All media was supplemented with 50 μg/mL penicillin–streptomycin (Gibco, ThermoFisher Scientific). BT and L3 cells were cultured as described previously ^26^.

### Immunoblots

Cells were harvested by NETN lysis as described previously ^27^. Antibodies to ATM (Upstate, #05-513), DNA-PKcs (generated in house), and Ku80 (Abcam #33242) were purchased as indicated.

### Generation of ATM-deficient cell lines

A549 cells with CRISPR/Cas9 deletion of ATM or DNA-PKcs have been described previously ^27^. Stable knock down of ATM in pancreatic cancer cells Panc 10.05 was achieved using shRNA as described previously ^20,36^. For CRISPR/Cas9 depletion of ATM in PC-3 cells, three short guide RNAs targeting exons 9, 16 and 23 of human ATM were designed, synthesized and ligated into pSpCas9(BB)-2A-GFP (pX458) vector, a gift from Feng Zhang (Addgene plasmid #48138). Constructs were transfected into PC-3 cells using Lipofectamine 2000 (Invitrogen) according to the manufacturer’s recommended procedures and GFP containing cells were sorted by flow cytometry. Genomic DNA was isolated from wild-type cells and PC-3 cells transfected with sgRNA constructs targeting ATM exons 9, 16 or 23. The DNA fragment around the target site was amplified by PCR and PCR products were cleaved by Surveyor Nuclease S as per the manufacturer’s recommended conditions. Surveyor Nuclease S assay confirmed the presence of CRISPR activity in exon 9, 16 and 23 sgRNA constructs. TIDE analysis showed that over 90% of cells transfected with sgRNA construct targeting on exon 16 contained insertion/deletion mutations. No cells were recovered from cells transfected with sgRNA construct targeting exon 9 (out of over 800 colonies screened) or targeting exon 23 (out of over 800 colonies screened). Only one clone, PC-3 E16 was recovered from the sgRNA construct targeting exon 16 (out of 2,400 colonies screened). Clone PC-3-E16 was expanded and DNA sequencing revealed that one allele of ATM had a one nucleotide (A) deletion in exon 16 which created a premature stop codon while the other allele had a 12-nucleotide deletion in exon 16 predicted to cause a one amino acid mutation and a four amino acid deletion. This clone was used for all subsequent experiments.

### Inhibitors

Olaparib and AZD6738 were purchased from Selleck Chemicals, dissolved in DMSO and stored as stock solutions at 10 mM at −20°C and −80°C respectively.

### Trypan blue viability assays

Trypan blue assays were carried out using trypan blue solution (Biorad) and a TC-20 automated cell counter (BioRad) according to the manufacturers’ instructions. Briefly, 50,000 to 100,000 cells were seeded for each experiment. Cells were allowed to attach for 16 hours, then treated with either 1 μM olaparib, 0.3 μM AZD6738, a combination of olaparib (1 µM) and AZD6738 (0.3 µM) or an equivalent volume of DMSO. Cells were harvested after 48, 96, 120 and 144 h. The mean with SEM of three independent experiments is shown. Statistical significance was determined by one-way ANOVA. P values of less than 0.05 were considered statistically significant and are indicated by an *.

### Annexin assays

Cells were seeded at 50,000 to 100,000 cells per 6 cm dish, treated as above and then analyzed by annexin staining and flow cytometry using an Annexin Apoptosis assay (ThermoFisher Scientific Cat#V13241). Cells were harvested after 48, 96, 120 and 144 h. The mean with SEM of three independent experiments is shown, and statistical significance was determined by one-way ANOVA. P values of less than 0.05 were considered statistically significant and are indicated by an *.

## Results

### ATM-deficient A549 lung cancer cells are sensitive to the combination of olaparib and AZD6738

ATM-deficient, DNA-PKcs-deficient, and control A549 cells were treated with olaparib (1 µM), AZD6738 (0.3 µM) or a combination of both inhibitors and viability was determined using the trypan blue exclusion assay. The viability of DNA-PKcs-deficient and control A549 cells was unaffected by the addition of either inhibitor alone or by a combination of both inhibitors (**Figure 1A, B**), although, as reported previously, A549-CRISPR-DNA-PKcs cells grew more slowly than control cells ^27,37^. Incubation with olaparib or AZD6738 alone decreased the viability of ATM-deficient cells by about 50% at the 114-hour time point, whereas incubation with both compounds decreased viability by about 90% (**Figure 1A, B).** As shown previously for VE-821 ^27^, neither olaparib nor AZD6738 induced apoptosis as single agents, but the combination of olaparib plus AZD6738 induced apoptosis in the ATM-deficient cells as determined by annexin staining, while leaving control ATM-proficient and DNA-PKcs-deficient cells unaffected (**Figure 1C**). Thus, similar to VE-821 ^27^, AZD6738 induces apoptosis in ATM-deficient A549 cells treated with olaparib, while ATM-competent cells are relatively unaffected by either agent alone or in combination (**Figure 1**).

**Figure 1:**
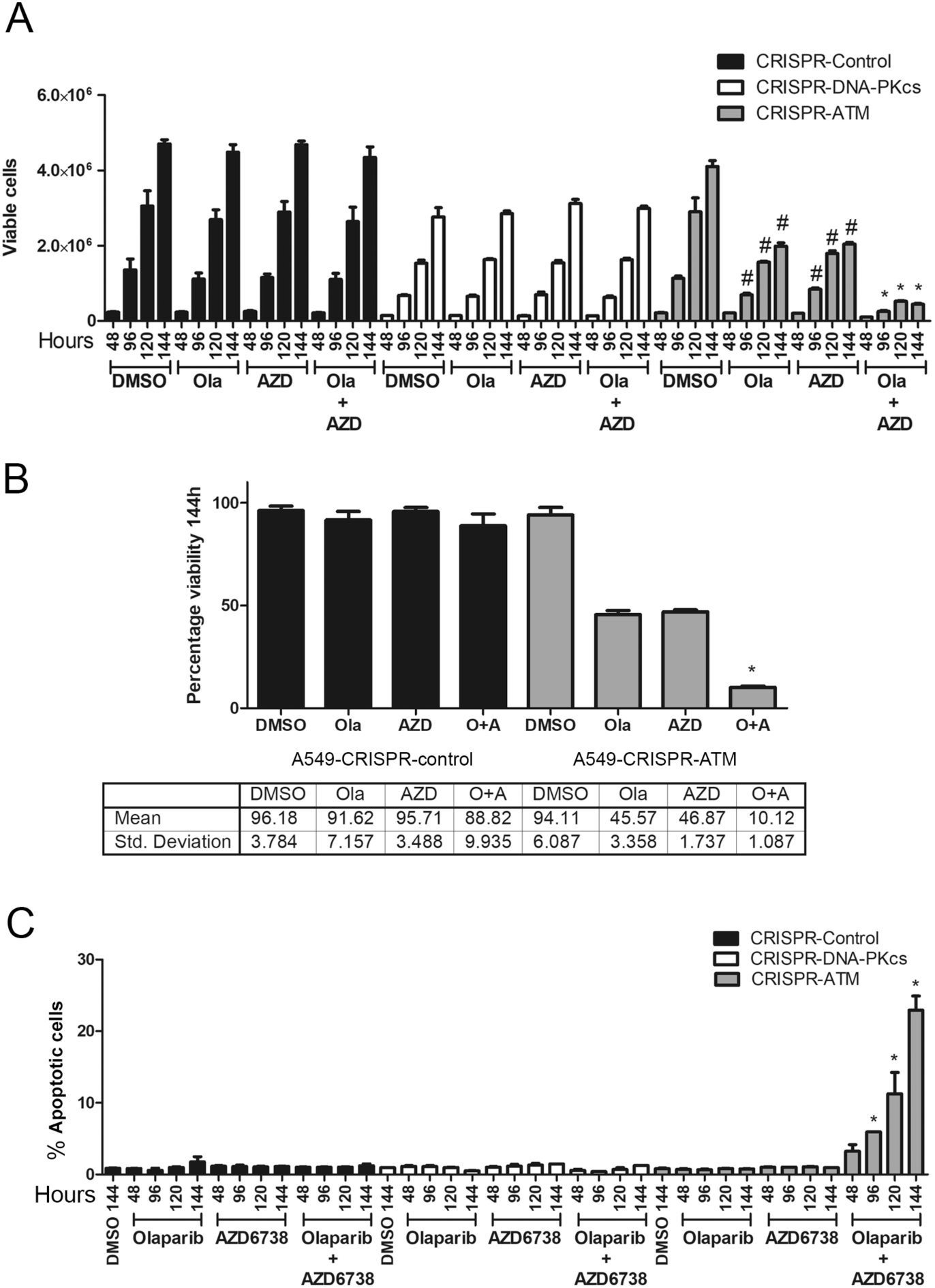
ATM-deficient A549 lung cancer cells undergo cell death when incubated with olaparib plus the ATR inhibitor AZD6738 but not either agent alone. A549-CRISPR-control (black bars), A549-CRISPR-DNA-PKcs (white bars) and A549-CRISPR-ATM cells (grey bars) generated as described ^27^ were incubated with olaparib (1 µM), AZD6738 (0.3 µM) or both olaparib (Ola) and AZD6738 (Azd) for 48-144 hours and assayed for either cell viability using the trypan blue exclusion assay (panel A) or apoptosis using annexin staining (panel C). Control samples were incubated with an equal volume of DMSO. Panel B shows quantitation of panel A at the 144-hour time point for ATM-proficient (left) and ATM-deficient (right) lung cancer cell lines. O represents olaparib and A represents AZD6738. Results show the mean with SEM of three separate experiments. Statistical significance was determined by one-way ANOVA. * Indicates p-value < 0.05 when compared to other treatment groups of that cell line and the DMSO control. In panel A, # indicates p-value < 0.05 when compared to DMSO control of that cell line only.

### ATM-deficient PC-3 prostate cancer cells are sensitive to the combination of olaparib and AZD6738

We next depleted ATM from PC-3 cells using CRISPR/Cas9 (**Figure 2A**). PC-3 cells are derived from bone metastasis of a prostate cancer patient and have homozygous *TP53* mutation. No mutations are reported for *ATM, BRCA1* or *2* or *PTEN* in the Catalogue of Somatic Mutations in Cancer (COSMIC) database ^38,39^, however PC-3 cells do not express PTEN protein^40^, possibly due to promoter methylation. Despite multiple attempts, we were unable to obtain complete deletion of ATM using CRISPR/Cas9, but we isolated one clone, E16-1, in which one allele of ATM had a one nucleotide (adenine) deletion in exon 16 which created a premature stop codon, while the other allele had a 12-nucleotide deletion in exon 16 predicted to cause a one amino acid mutation and a four amino acid deletion. Western blot indicated that this clone expressed very low levels of ATM protein, approximately 10% of that in control PC-3 cells (**Figure 2A)**. ATM-deficient and control PC-3 cells were treated with olaparib (1 µM), AZD6738 (0.3 µM) or a combination of both inhibitors and assayed for viability and cell death as above. The decrease in viability at the 144-hour time point in the ATM-deficient cell lines was 28% and 39% for olaparib and AZD6738 alone, respectively, and 83% for the combination of olaparib plus AZD6738 (**Figure 2B, C**). Importantly, cell death, as determined by the annexin staining, was only observed in ATM-deficient PC-3 cells treated with both olaparib and AZD6738 (**Figure 2D**).

**Figure 2:**
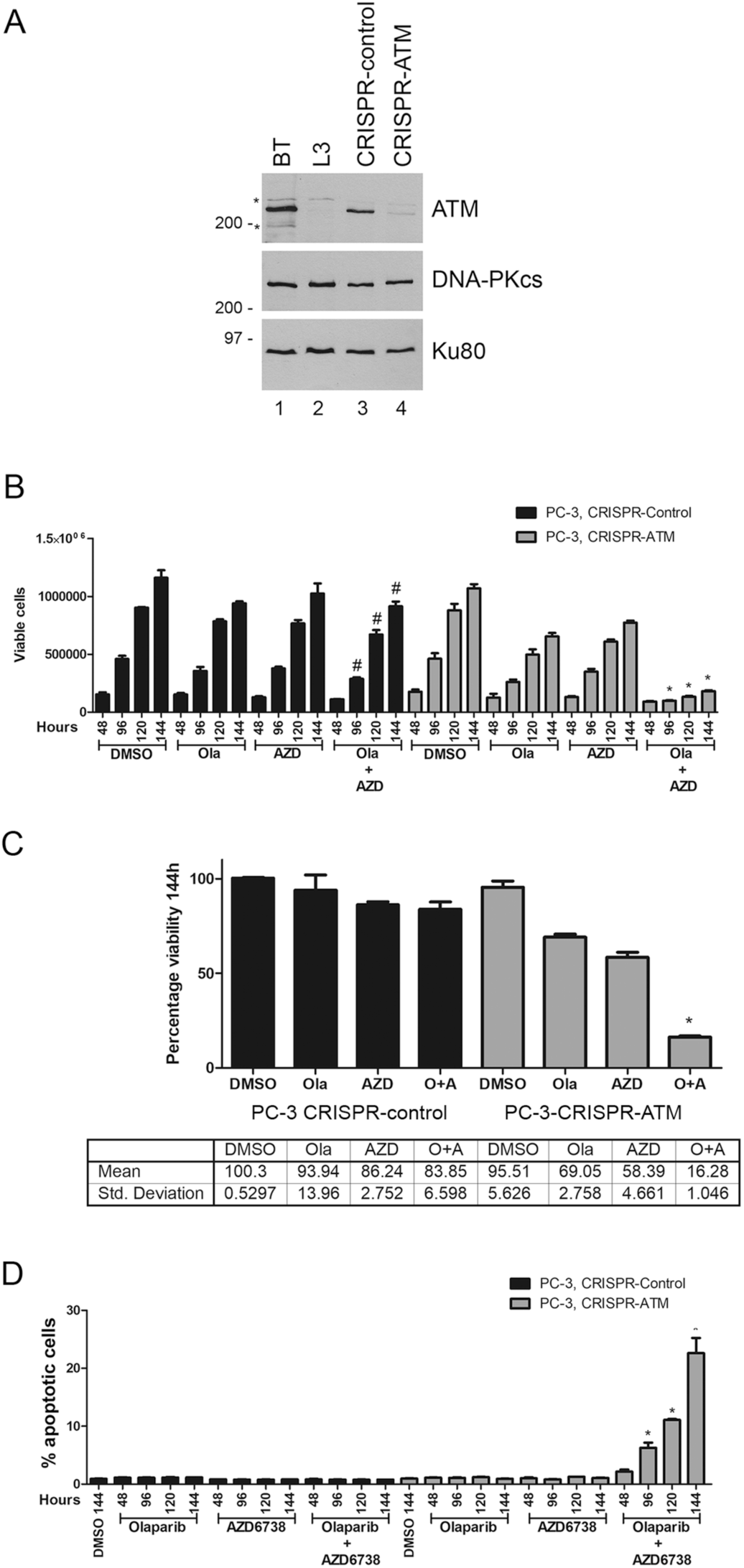
ATM-deficient PC-3 prostate cancer cells undergo cell death when incubated with olaparib plus the ATR inhibitor AZD6738 but not either agent alone. Panel A, ATM was depleted from PC-3 prostate cancer cells using CRISPR/Cas9 as described above. Extracts were generated by NETN lysis and 50 µg total protein run on SDS PAGE and immunoblotted for ATM, DNA-PKcs or Ku80 as shown (panel A). BT and L3 are lymphoblastoid cell lines from a control (BT) and an A-T patient (L3) as described previously ^20^. Molecular weight markers are indicated in kDa on the right-hand side. * represents non-specific cross-reacting bands. The effects of olaparib and AZD6738 on cell viability and apoptosis are shown in panels B and C (viability) and D (apoptosis). * indicates p-value < 0.05 when compared to other treatment groups of that cell line and the DMSO control. # indicates p-value < 0.05 when compared to DMSO control of that cell line only. Panel C shows quantitation of panel A at the 144-hour time point for ATM-proficient (left) and ATM-deficient (right) prostate cancer cell lines.

### ATM-deficient Panc-10.05 pancreatic cancer cells are sensitive to the combination of olaparib and AZD6738

To test whether ATM-deficient pancreatic cancer cells are also sensitive to olaparib and AZD6738, we depleted ATM from Panc 10.05 cells using shRNA, as described previously ^20,22,26^. Knock-down of ATM is shown by western blot in **Figure 3A**. Panc 10.05 cells are heterozygous for c.3077+103A>G mutation in *ATM* and homozygous for I255N in *TP53*. ATM-deficient and shGFP Panc 10.05 (control) cells were treated with olaparib (1 µM), AZD6738 (0.3 µM) or a combination of both inhibitors as above. As in lung and prostate cancer cells, olaparib and AZD6738 alone had no significant effect on viability in ATM-proficient pancreatic cancer cells, whereas the combination of olaparib plus AZD6738 reduced the number of viable cells by almost 70% compared to 6 and 17% for either treatment alone (**Figure 3B, C**). Moreover, as in lung and prostate cancer cells, cell death was only observed in ATM-deficient cells treated with the combination of olaparib plus AZD6738 and no cell death was observed in ATM-proficient pancreatic cancer cells treated with olaparib and AZD6738 alone or in combination (**Figure 3D**).

**Figure 3:**
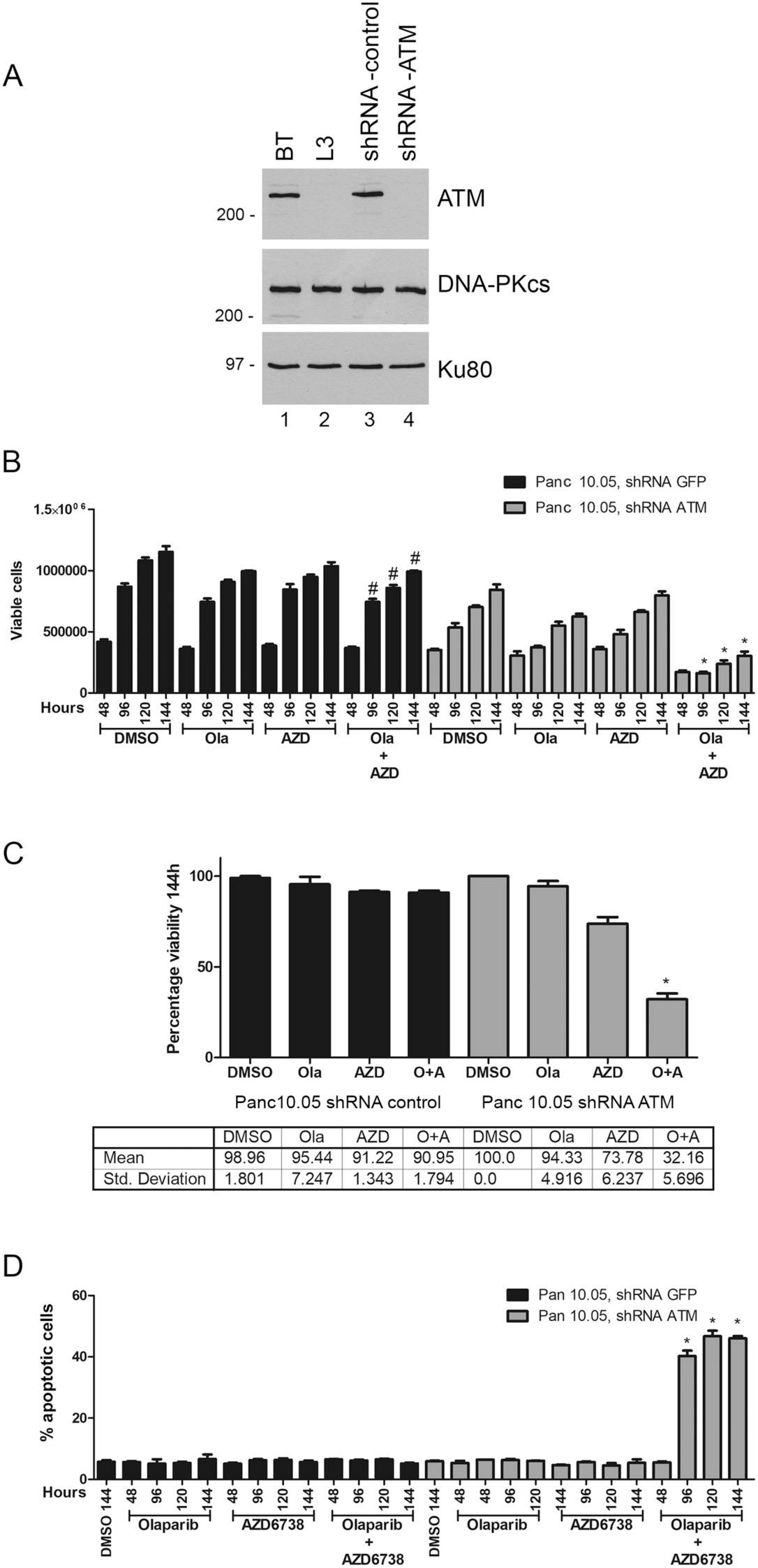
ATM-deficient Panc 10.05 pancreatic cancer cells undergo cell death when incubated with olaparib plus the ATR inhibitor AZD6738 but not either agent alone. Panel A, ATM was depleted from Panc 10.05 pancreatic cancer cells using shRNA as described previously^20^. Extracts were generated by NETN lysis and 50 µg total protein run on SDS PAGE and immunoblotted for ATM, DNA-PKcs or Ku80 as shown. BT and L3 are ATM positive and negative controls as in Figure 2. The effect of olaparib and AZD6738 on viability and apoptosis are shown in panels B and C (viability) and D (apoptosis). Quantitation of viability at the 114-hour time point is shown in panel C. Statistical analysis and * are as in Figures 1 and 2.

## Discussion

PARP inhibitors are proving to be highly effective in targeting cancers with mutations in the *BRCA1* and *BRCA2* genes. Since BRCA1 and BRCA2 proteins play key roles in the detection and repair of DSBs, these findings open up the possibility that cancers with mutations in other DNA repair genes may also respond to PARP inhibitor therapy. One such candidate is ATM, a DNA damage activated protein kinase that plays an important role in phosphorylation of both p53, and checkpoint kinase 2 (Chk2), and regulation of cell cycle checkpoints and cell survival after DNA damage, as well as repair of complex DNA DSBs ^41^. Moreover, the *ATM* gene is mutated in a number of cancers including lymphoma, lung, prostate and pancreatic cancers ^42,43^, offering opportunities for precision therapy.

We previously showed that the PARP inhibitor olaparib reduces cell proliferation and induces reversible G2 arrest in ATM-deficient A549 lung cancer cells ^27^. We hypothesized that disruption of the G2 checkpoint using an ATR inhibitor would overcome the olaparib-induced G2/M arrest and induce cell death. Indeed, we found that the combination of the ATR inhibitor VE-821 with olaparib induced apoptosis in ATM-depleted A549 lung cancer cells with little effect on viability of ATM-proficient A549 cells ^27^. Here, we extend these studies to show that the ATR inhibitor AZD6738 also induces cell death in olaparib-treated ATM-deficient lung, prostate and pancreatic cancer cells with little effect on ATM-proficient control cells. This suggests that the combination of a PARP inhibitor with an ATR inhibitor may be useful in the treatment of a variety of cancer types with ATM-deficiency. Our findings may also have benefit in cases of PARP inhibitor resistance, and/or in patients with mutations in other DDR gene subsets that make tumours less responsive to PARP inhibition. In each case, we speculate that the combination of PARP inhibitor with ATR inhibitor could have benefit over either agent alone.

As discussed, PARP inhibitors are FDA-approved for use in ovarian cancer and are in clinical trials for a number of other cancers, including breast, prostate and pancreatic cancer. In an initial clinical trial of 60 patients with metastatic prostate cancer, five were shown to have somatic or germline mutations in *ATM* and 4 of the 5 responded favourably to olaparib ^32^, suggesting that ATM is a promising target in prostate cancer. Subsequently, olaparib treatment was shown to result in statistically significant increase in radiographic progression free survival (rPFS) compared to physician’s choice therapy in patients with metastatic castration resistant prostate cancer and DDR gene alterations, for example in *BRCA1/2, ATM* and *PALB2* ^44^. However, exploratory analyses suggest that patients with *ATM* gene defects may not experience as impressive rates of progression-free survival in response to olaparib treatment alone as patients with other DDR gene alterations. This early evidence supports the need to examine combination therapy to enhance the clinical efficacy in this unique DDR gene alteration. Our observation that olaparib alone does not cause cell death in ATM-deficient prostate cancer cell lines, may explain why men with mutation in *ATM* exhibited inferior outcome to olaparib treatment compared to those with BRCA1 mutation ^45^, supporting our hypothesis that combination of PARP inhibitor with an ATR inhibitor may be more appropriate for tumours with ATM-deficiency.

Critical to adoption of PARP inhibitors in the clinic will be the accurate identification of patients who may benefit from the selected treatment. *ATM* mutations in patient cancers are mainly mis-sense mutations and are scattered throughout the coding sequence ^42,43^. Only R337 in the amino terminus and K2811 and R3008 in the C-terminal kinase domain recur in multiple tumours and may represent more frequent mutations. Moreover, the functional consequences of these mutations are unknown, thus screening by DNA sequence alone may not be informative. However, analysis of ATM mutations in Ataxia Telangiectasia, a cancer predisposition syndrome caused by loss or inactivation of both copies of the ATM gene ^46^ has shown that many mis-sense mutations result in protein truncation ^47^. Indeed, immunohistochemistry of ATM protein expression in lung adenocarcinoma revealed that over 40% had low protein expression, despite only 10% showing gene mutation ^48^. In addition, *ATM* promoter methylation has been shown to lead to silencing in some cancer cells ^49,50^. Therefore, it is possible that more patient samples may lack ATM expression or function than indicated by exome sequencing and that methods for assaying ATM protein expression and/or activity will be required for patient selection.

Our results also highlight how the type of assay used to measure sensitivity of cultured cells to a compound can influence interpretation of results. Our results using clonogenic survival assays indicated that ATM-deficient cells are highly sensitive to olaparib (less than 1% survival at 1 µM olaparib ^27^), however, subsequent results based on a better understanding of the mechanism of action of olaparib, indicated that these cells were only arrested in G2 and did not show evidence of cell death by either sub-G1 DNA or annexin staining. Only after including the ATR inhibitor VE-821 did we observed evidence of cell death ^27^. These data illustrate how inferring cell viability from a clonogenic survival/colony formation assay^51^ can be misleading, as the assay measures the ability of a single cell to proliferate to form a colony, usually of 50 or more cells, not viability/cell death, and, as shown by our work, a cell that is arrested in G2 is still viable; it is just unable to form colonies. Therefore, the type of viability assay used for determining sensitivity to olaparib and possibly other PARP inhibitors, needs to be carefully chosen.

## Authorship

Experiments and data analysis were carried out by SR, NRJ, RY, YY, GA, MKu, SG and SPLM. The study was conceived by MK and SPLM with intellectual contributions from PB, SY and TAB. All authors contributed to writing and editing of the manuscript and have approved the final submission.

## Acknowledgements

We thank the Flow Cytometry Facility and the Center for Genome Engineering at the Cumming School of Medicine, University of Calgary for expert assistance.

## References

1. Pommier Y, O’Connor MJ, de Bono J. Laying a trap to kill cancer cells: PARP inhibitors and their mechanisms of action. Sci Transl Med 2016; 8(362): 362ps317; e-pub ahead of print 2016/11/01; doi 10.1126/scitranslmed.aaf9246.

2. Bryant HE, Schultz N, Thomas HD, Parker KM, Flower D, Lopez E et al. Specific killing of BRCA2-deficient tumours with inhibitors of poly(ADP-ribose) polymerase. Nature 2005; 434(7035): 913-917; e-pub ahead of print 2005/04/15; doi 10.1038/nature03443.

3. Farmer H, McCabe N, Lord CJ, Tutt AN, Johnson DA, Richardson TB et al. Targeting the DNA repair defect in BRCA mutant cells as a therapeutic strategy. Nature 2005; 434(7035): 917-921; e-pub ahead of print 2005/04/15; doi 10.1038/nature03445.

4. FDA approved olaparib (LYNPARZA, AstraZeneca Pharmaceuticals LP) for the maintenance treatment of adult patients with deleterious or suspected deleterious germline or somatic BRCA-mutated (gBRCAm or sBRCAm) advanced epithelial ovarian, fallopian tube or primary peritoneal cancer who are in complete or partial response to first-line platinum-based. https://www.fda.gov/drugs/fda-approved-olaparib-lynparza-astrazeneca-pharmaceuticals-lp-maintenance-treatment-adult-patients (2018).

5. Lord CJ, Ashworth A. PARP inhibitors: Synthetic lethality in the clinic. Science (New York, NY) 2017; 355(6330): 1152-1158; e-pub ahead of print 2017/03/18; doi10.1126/science.aam7344.

6. Coleman RL, Oza AM, Lorusso D, Aghajanian C, Oaknin A, Dean A et al. Rucaparib maintenance treatment for recurrent ovarian carcinoma after response to platinum therapy (ARIEL3): a randomised, double-blind, placebo-controlled, phase 3 trial. Lancet 2017; 390(10106): 1949-1961; e-pub ahead of print 2017/09/17; doi10.1016/s0140-6736(17)32440-6.

7. Golan T, Hammel P, Reni M, Van Cutsem E, Macarulla T, Hall MJ et al. Maintenance Olaparib for Germline BRCA-Mutated Metastatic Pancreatic Cancer. The New England journal of medicine 2019; 381(4): 317-327; e-pub ahead of print 2019/06/04; doi10.1056/NEJMoa1903387.

8. Mateo J, Porta N, Bianchini D, McGovern U, Elliott T, Jones R et al. Olaparib in patients with metastatic castration-resistant prostate cancer with DNA repair gene aberrations (TOPARP-B): a multicentre, open-label, randomised, phase 2 trial. The Lancet Oncology 2019; e-pub ahead of print 2019/12/07; doi10.1016/s1470-2045(19)30684-9.

9. Mirza MR, Monk BJ, Herrstedt J, Oza AM, Mahner S, Redondo A et al. Niraparib Maintenance Therapy in Platinum-Sensitive, Recurrent Ovarian Cancer. The New England journal of medicine 2016; 375(22): 2154-2164; e-pub ahead of print 2016/10/09; doi10.1056/NEJMoa1611310.

10. Moore K, Colombo N, Scambia G, Kim BG, Oaknin A, Friedlander M et al. Maintenance Olaparib in Patients with Newly Diagnosed Advanced Ovarian Cancer. The New England journal of medicine 2018; 379(26): 2495-2505; e-pub ahead of print 2018/10/23; doi10.1056/NEJMoa1810858.

11. Robson M, Im SA, Senkus E, Xu B, Domchek SM, Masuda N et al. Olaparib for Metastatic Breast Cancer in Patients with a Germline BRCA Mutation. The New England journal of medicine 2017; 377(6): 523-533; e-pub ahead of print 2017/06/06; doi10.1056/NEJMoa1706450.

12. Mateo J, Carreira S, Sandhu S, Miranda S, Mossop H, Perez-Lopez R et al. DNA-Repair Defects and Olaparib in Metastatic Prostate Cancer. The New England journal of medicine 2015; 373(18): 1697-1708; e-pub ahead of print 2015/10/29; doi 10.1056/NEJMoa1506859.

13. Boulton SJ. Cellular functions of the BRCA tumour-suppressor proteins. Biochemical Society transactions 2006; 34(Pt 5): 633-645; e-pub ahead of print 2006/10/21; doi 10.1042/bst0340633.

14. McCabe N, Turner NC, Lord CJ, Kluzek K, Bialkowska A, Swift S et al. Deficiency in the repair of DNA damage by homologous recombination and sensitivity to poly(ADP-ribose) polymerase inhibition. Cancer research 2006; 66(16): 8109-8115; e-pub ahead of print 2006/08/17; doi 66/16/8109 [pii] 10.1158/0008-5472.CAN-06-0140.

15. Bryant HE, Helleday T. Inhibition of poly (ADP-ribose) polymerase activates ATM which is required for subsequent homologous recombination repair. Nucleic acids research 2006; 34(6): 1685–1691.

16. Shiloh Y. ATM: Expanding roles as a chief guardian of genome stability. Experimental cell research 2014; e-pub ahead of print 2014/09/15; doi 10.1016/j.yexcr.2014.09.002.

17. Shiloh Y, Lederman HM. Ataxia-telangiectasia (A-T): An emerging dimension of premature ageing. Ageing research reviews 2016; e-pub ahead of print 2016/05/18; doi 10.1016/j.arr.2016.05.002.

18. Choi M, Kipps T, Kurzrock R. ATM Mutations in Cancer: Therapeutic Implications. Molecular cancer therapeutics 2016; 15(8): 1781–1791; e-pub ahead of print 2016/07/15; doi 10.1158/1535-7163.mct-15-0945.

19. Greiner TC, Dasgupta C, Ho VV, Weisenburger DD, Smith LM, Lynch JC et al. Mutation and genomic deletion status of ataxia telangiectasia mutated (ATM) and p53 confer specific gene expression profiles in mantle cell lymphoma. Proceedings of the National Academy of Sciences of the United States of America 2006; 103(7): 2352–2357; e-pub ahead of print 2006/02/08; doi 10.1073/pnas.0510441103.

20. Williamson CT, Muzik H, Turhan AG, Zamo A, O’Connor MJ, Bebb DG et al. ATM deficiency sensitizes mantle cell lymphoma cells to poly(ADP-ribose) polymerase-1 inhibitors. Molecular cancer therapeutics 2010; 9(2): 347–357; e-pub ahead of print 2010/02/04; doi 10.1158/1535-7163.mct-09-0872.

21. Weston VJ, Oldreive CE, Skowronska A, Oscier DG, Pratt G, Dyer MJ et al. The PARP inhibitor olaparib induces significant killing of ATM-deficient lymphoid tumor cells in vitro and in vivo. Blood 2010; 116(22): 4578–4587; e-pub ahead of print 2010/08/27; doi 10.1182/blood-2010-01-265769.

22. Williamson CT, Kubota E, Hamill JD, Klimowicz A, Ye R, Muzik H et al. Enhanced cytotoxicity of PARP inhibition in mantle cell lymphoma harbouring mutations in both ATM and p53. EMBO molecular medicine 2012; 4(6): 515–527; e-pub ahead of print 2012/03/15; doi 10.1002/emmm.201200229.

23. Kim HS, Choi SI, Min HL, Kim MA, Kim WH. Mutation at intronic repeats of the ataxia-telangiectasia mutated (ATM) gene and ATM protein loss in primary gastric cancer with microsatellite instability. PloS one 2013; 8(12): e82769; e-pub ahead of print 2013/12/11; doi 10.1371/journal.pone.0082769.

24. Kim HS, Kim MA, Hodgson D, Harbron C, Wellings R, O’Connor MJ et al. Concordance of ATM (ataxia telangiectasia mutated) immunohistochemistry between biopsy or metastatic tumor samples and primary tumors in gastric cancer patients. Pathobiology 2013; 80(3): 127–137; e-pub ahead of print 2013/01/19; doi 10.1159/000346034.

25. Kubota E, Williamson CT, Ye R, Elegbede A, Peterson L, Lees-Miller SP et al. Low ATM protein expression and depletion of p53 correlates with olaparib sensitivity in gastric cancer cell lines. Cell cycle (Georgetown, Tex) 2014; 13(13): 2129–2137; e-pub ahead of print 2014/05/21; doi 10.4161/cc.29212.

26. Wang C, Jette N, Moussienko D, Bebb DG, Lees-Miller SP. ATM-Deficient Colorectal Cancer Cells Are Sensitive to the PARP Inhibitor Olaparib. Transl Oncol 2017; 10(2): 190–196; e-pub ahead of print 2017/02/10; doi 10.1016/j.tranon.2017.01.007.

27. Jette NR, Radhamani S, Arthur G, Ye R, Goutam S, Bolyos A et al. Combined poly-ADP ribose polymerase and ataxia-telangiectasia mutated/Rad3-related inhibition targets ataxia-telangiectasia mutated-deficient lung cancer cells. British journal of cancer 2019; 121(7): 600–610; e-pub ahead of print 2019/09/05; doi 10.1038/s41416-019-0565-8.

28. Reaper PM, Griffiths MR, Long JM, Charrier JD, Maccormick S, Charlton PA et al. Selective killing of ATM- or p53-deficient cancer cells through inhibition of ATR. Nat Chem Biol 2011; 7(7): 428–430; e-pub ahead of print 2011/04/15; doi 10.1038/nchembio.573.

29. Foote KM, Lau A, Nissink JW. Drugging ATR: progress in the development of specific inhibitors for the treatment of cancer. Future medicinal chemistry 2015; 7(7): 873–891; e-pub ahead of print 2015/06/11; doi 10.4155/fmc.15.33.

30. Foote KM, Nissink JWM, McGuire T, Turner P, Guichard S, Yates JWT et al. Discovery and Characterization of AZD 6738, a Potent Inhibitor of Ataxia Telangiectasia Mutated and Rad3 Related (ATR) Kinase with Application as an Anticancer Agent. J Med Chem 2018; 61(22): 9889–9907; e-pub ahead of print 2018/10/23; doi 10.1021/acs.jmedchem.8b01187.

31. Canadian Cancer Statistics 2019. http://www.cancer.ca/statistics:

32. Mateo J, Boysen G, Barbieri CE, Bryant HE, Castro E, Nelson PS et al. DNA Repair in Prostate Cancer: Biology and Clinical Implications. Eur Urol 2017; 71(3): 417–425; e-pub ahead of print 2016/10/30; doi 10.1016/j.eururo.2016.08.037.

33. Grant RC, Selander I, Connor AA, Selvarajah S, Borgida A, Briollais L et al. Prevalence of Germline Mutations in Cancer Predisposition Genes in Patients with Pancreatic Cancer. Gastroenterology 2014; e-pub ahead of print 2014/12/06; doi 10.1053/j.gastro.2014.11.042.

34. Roberts NJ, Jiao Y, Yu J, Kopelovich L, Petersen GM, Bondy ML et al. ATM mutations in patients with hereditary pancreatic cancer. Cancer discovery 2012; 2(1): 41–46; e-pub ahead of print 2012/05/16; doi 10.1158/2159-8290.cd-11-0194.

35. Kim H, Saka B, Knight S, Borges M, Childs E, Klein A et al. Having pancreatic cancer with tumoral loss of ATM and normal TP53 protein expression is associated with a poorer prognosis. Clinical cancer research : an official journal of the American Association for Cancer Research 2014; 20(7): 1865–1872; e-pub ahead of print 2014/02/04; doi 10.1158/1078-0432.ccr-13-1239.

36. Elkon R, Rashi-Elkeles S, Lerenthal Y, Linhart C, Tenne T, Amariglio N et al. Dissection of a DNA-damage-induced transcriptional network using a combination of microarrays, RNA interference and computational promoter analysis. Genome biology 2005; 6(5): R43.

37. Ruis BL, Fattah KR, Hendrickson EA. The catalytic subunit of DNA-dependent protein kinase regulates proliferation, telomere length, and genomic stability in human somatic cells. Molecular and cellular biology 2008; 28(20): 6182–6195.

38. Bamford S, Dawson E, Forbes S, Clements J, Pettett R, Dogan A et al. The COSMIC (Catalogue of Somatic Mutations in Cancer) database and website. British journal of cancer 2004; 91(2): 355–358; e-pub ahead of print 2004/06/10; doi 10.1038/sj.bjc.6601894 6601894 [pii].

39. Forbes SA, Bindal N, Bamford S, Cole C, Kok CY, Beare D et al. COSMIC: mining complete cancer genomes in the Catalogue of Somatic Mutations in Cancer. Nucleic acids research 2011; 39(Database issue): D945–950; e-pub ahead of print 2010/10/19; doi gkq929 [pii] 10.1093/nar/gkq929.

40. Fraser M, Zhao H, Luoto KR, Lundin C, Coackley C, Chan N et al. PTEN deletion in prostate cancer cells does not associate with loss of RAD51 function: implications for radiotherapy and chemotherapy. Clinical cancer research : an official journal of the American Association for Cancer Research 2012; 18(4): 1015–1027; e-pub ahead of print 2011/11/25; doi 10.1158/1078-0432.Ccr-11-2189.

41. Blackford AN, Jackson SP. ATM, ATR, and DNA-PK: The Trinity at the Heart of the DNA Damage Response. Molecular cell 2017; 66(6): 801–817; e-pub ahead of print 2017/06/18; doi 10.1016/j.molcel.2017.05.015.

42. Gao J, Aksoy BA, Dogrusoz U, Dresdner G, Gross B, Sumer SO et al. Integrative analysis of complex cancer genomics and clinical profiles using the cBioPortal. Science signaling 2013; 6(269): pl1; e-pub ahead of print 2013/04/04; doi 10.1126/scisignal.2004088.

43. Cerami E, Gao J, Dogrusoz U, Gross BE, Sumer SO, Aksoy BA et al. The cBio cancer genomics portal: an open platform for exploring multidimensional cancer genomics data. Cancer discovery 2012; 2(5): 401–404; e-pub ahead of print 2012/05/17; doi 10.1158/2159-8290.cd-12-0095.

44. Targeted Therapy Slows Progression of Advanced Prostate Cancer [ESMO 2019 Press Release]. 2019.

45. Marshall CH, Sokolova AO, McNatty AL, Cheng HH, Eisenberger MA, Bryce AH et al. Differential Response to Olaparib Treatment Among Men with Metastatic Castration-resistant Prostate Cancer Harboring BRCA1 or BRCA2 Versus ATM Mutations. Eur Urol 2019; 76(4): 452–458; e-pub ahead of print 2019/02/25; doi 10.1016/j.eururo.2019.02.002.

46. Rothblum-Oviatt C, Wright J, Lefton-Greif MA, McGrath-Morrow SA, Crawford TO, Lederman HM. Ataxia telangiectasia: a review. Orphanet journal of rare diseases 2016; 11(1): 159; e-pub ahead of print 2016/11/26; doi 10.1186/s13023-016-0543-7.

47. Gilad S, Chessa L, Khosravi R, Russell P, Galanty Y, Piane M et al. Genotype-phenotype relationships in ataxia-telangiectasia and variants. Am J Hum Genet 1998; 62(3): 551–561.

48. Villaruz LC, Jones H, Dacic S, Abberbock S, Kurland BF, Stabile LP et al. ATM protein is deficient in over 40% of lung adenocarcinomas. Oncotarget 2016; e-pub ahead of print 2016/06/04; doi 10.18632/oncotarget.9757.

49. Hirakawa T, Nasu K, Aoyagi Y, Takebayashi K, Zhu R, Narahara H. ATM expression is attenuated by promoter hypermethylation in human ovarian endometriotic stromal cells. Molecular human reproduction 2019; 25(6): 295–304; e-pub ahead of print 2019/03/15; doi 10.1093/molehr/gaz016.

50. Kim WJ, Vo QN, Shrivastav M, Lataxes TA, Brown KD. Aberrant methylation of the ATM promoter correlates with increased radiosensitivity in a human colorectal tumor cell line. Oncogene 2002; 21(24): 3864–3871.

51. Franken NA, Rodermond HM, Stap J, Haveman J, van Bree C. Clonogenic assay of cells in vitro. Nat Protoc 2006; 1(5): 2315–2319; e-pub ahead of print 2007/04/05; doi 10.1038/nprot.2006.339.

